# Genome-wide diversity in temporal and regional populations of the betabaculovirus *Erinnyis ello granulovirus* (ErelGV)

**DOI:** 10.1101/273672

**Authors:** A. F. Brito, F. L. Melo, D. M. P Ardisson-Araújo, W. Sihler, M. L. Souza, B.M. Ribeiro

## Abstract

**Background:** *Erinnyis ello granulovirus* (ErelGV) is a betabaculovirus infecting caterpillars of the sphingid moth *E. ello ello* (cassava hornworm), an important pest of cassava crops (*Manihot esculenta*). In this study, the genome of seven field isolates of the virus ErelGV were deep sequenced and their inter-and intrapopulational sequence diversity were analyzed.

**Results:** No events of gene gain/loss or translocations were observed, and indels were mainly found within highly repetitive regions (direct repeats, *drs*). A naturally occurring isolate from Northern Brazil (Acre State, an Amazonian region) has shown to be the most diverse population, with a unique pattern of polymorphisms. Overall, non-synonymous substitutions were found all over the seven genomes, with no specific gathering of mutations on hotspot regions. Independently of their sizes, some ORFs have shown higher levels of non-synonymous changes than others. Non-core genes of known functions and structural genes were among the most diverse ones; and as expected, core genes were the least variable genes. We observed remarkable differences on diversity of paralogous genes, as in multiple copies of *p10, fgf*, and *pep*. Another important contrast on sequence diversity was found on genes encoding complex subunits and/or involved in the same biological processes, as *late expression factors* (*lefs*) and *per os infectivity factors* (*pifs*). Interestingly, several polymorphisms in coding regions lie on sequences encoding specific protein domains.

**Conclusions:** By comparing and integrating information about inter-and intrapopulational diversity of viral isolates, we provide a detailed description on how evolution operates on field isolates of a betabaculovirus. Our results revealed that 35-41% of the SNPs of ErelGV lead to amino acid changes (non-synonymous substitutions). Some genes, especially non-core genes of unknown functions, tend to accumulate more mutations, while core genes evolve slowly and are more conserved. Additional studies would be necessary to understand the actual effects of such gene variations on viral infection and fitness.

## BACKGROUND

Baculoviruses are double-stranded DNA viruses found infecting insects from three different orders [1]. The *Baculoviridae* family is divided into four genera [2], the lepidopteran-specific (*Alphabaculovirus* and *Betabaculovirus*), hymenopteran-specific (*Gammabaculovirus*), and dipteran-specific viruses (*Deltabaculovirus*). Some of these viruses are used as efficient and sustainable bioinsecticides for controlling populations of pests in forests and crops, and are safe alternatives to chemical pesticides [3, 4]. Two of the best examples of baculoviral species used as insecticides are *Anticarsia gemmatalis multiple nucleopolyhedrovirus* (AgMNPV, an alphabaculovirus), in soybean crops [3, 5], and *Cydia pomonella granulovirus* (CpGV, a betabaculovirus), in fruit crops [6].

*Erinnyis ello* (Lepidoptera: Sphingidae) is a serious pest of cassava (*Manihot esculenta*) in the neotropics, with a broad geographic range extending from southern Brazil, Argentina, and Paraguay to the Caribbean basin and the southern United States [7]. This insect is also a severe pest of rubber tree (*Hevea* b*rasiliensis* M. Arg.). Several natural enemies of this insect have been identified including parasites, predators, fungi, bacteria, and a virus (*Erinnyis ello granulovirus*). Because of the migratory behavior of hornworm adults, this abundance of natural enemies does not prevent periodic caterpillar outbreaks [8].

In Brazil, the *Erinnyis ello granulovirus* (ErelGV) has been used efficiently to control populations of the cassava hornworm (*Erinnyis ello ello*) in cassava plantations [9, 10] (Figure 1) and very few studies on the biology and molecular characterization of this virus have been carried out [11, 12]. Cassava is a native plant from Brazil and an important carbohydrate source for human consumption, especially in Africa, Asia, and Latin America [13]. Given the agricultural importance of cassava, and the economic impacts of *E. ello* caterpillars, a recent study have characterized the virus morphology, genome sequence, and evolutionary history of ErelGV [11]. It revealed a 102,759 bp genome that lacks typical *homologous regions* (*hrs*) and encodes at least 130 ORFs. As for all baculoviruses, ErelGV also encodes a set of 38 genes shared by all baculoviruses, called ‘core genes’ [14]. Recently, Ardisson-Araujo et al. [12] constructed a recombinant Autographa californica multiple nucleopolyhedrovirus (AcMNPV) containing the the *tmk-dut* fusion gene (*erel5*) of ErelGV showing that the recombinant virus was able to accelerated viral DNA replication, BV and OB production.

**Figure 1.**
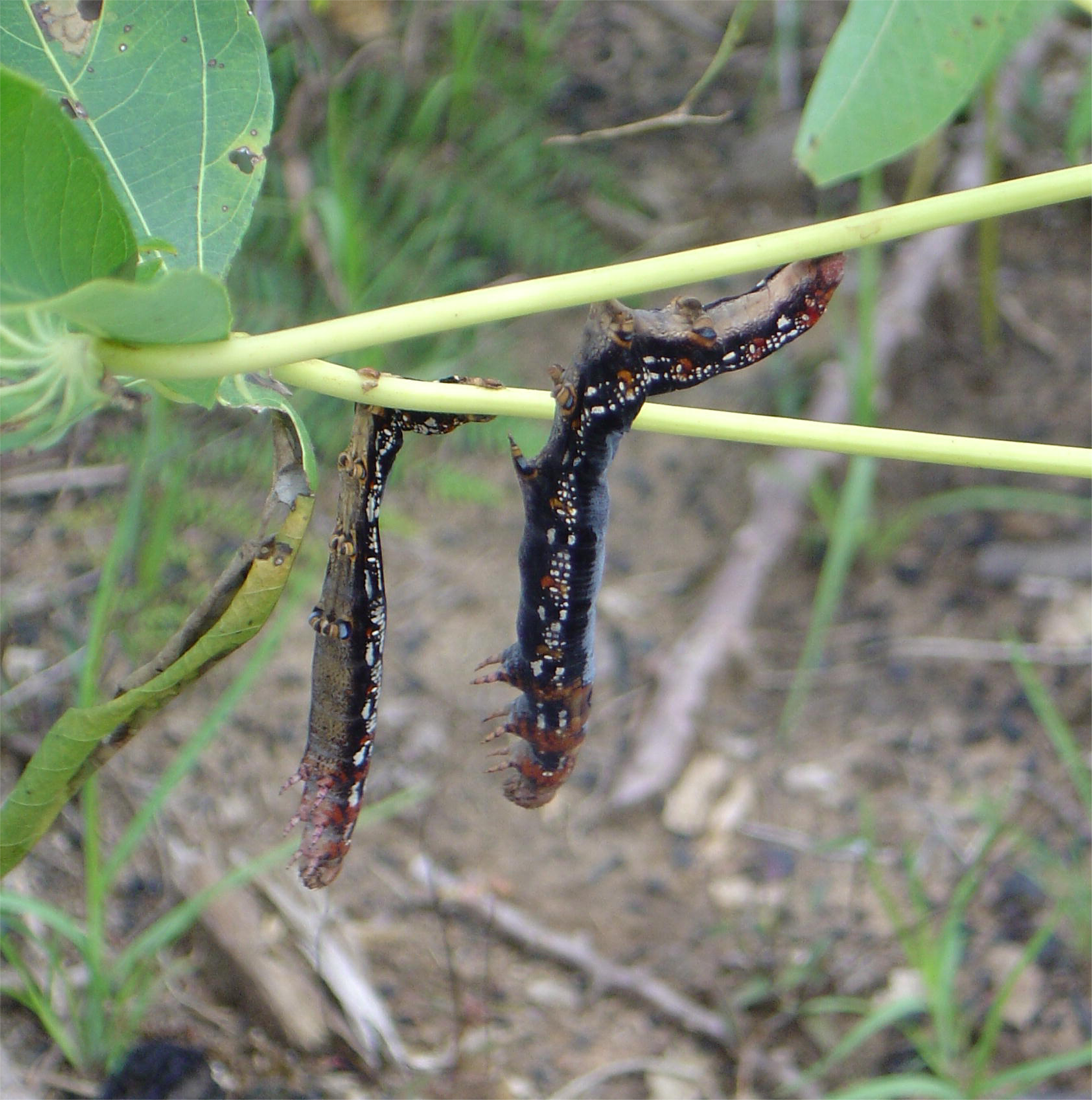
Caterpillars of *Erinnyis ello ello* (Lepidoptera: Sphingidae) infected by ErelGV in a Cassava crop in Acre State (AC), Brazil. Photo courtesy of Dr Murilo Fazolin.

Field populations of betabaculoviruses are known to be composed by multiple genotypic variants [15], however, few studies on genome sequence diversity of baculovirus isolates have been done [16]. Inter-isolate comparative studies have shown that baculoviral genes have low levels of polymorphisms, and non-synonymous substitutions (NSS) tend to be located within highly variable genes or specific genomic regions [17-19]. Gain and loss of genomic fragments are observed especially in repetitive regions, such as *homologous regions* (*hrs*) and *direct repeats* (*drs*) [15-17].

In this study, we investigated the inter-and intra-isolate genetic variability of seven temporal and regional field populations of ErelGV. Aspects of the ErelGV genomic organization and evolution are discussed; and we offer a detailed summary of polymorphisms on genes belonging to different functional categories.

## MATERIALS AND METHODS

### Viral samples and granules purification

The samples “ErelGV-86”, “-94”, “-98”, “-99” and “-00” correspond to Brazilian field isolates of ErelGV sequentially collected in 1986, 1994, 1998, 1999 and 2000 in cassava crops in Itajaí/Jaguaruna region (Santa Catarina State, Brazil), where these viruses were used for controlling populations of *E. ello ello* caterpillars (Figure 2). The “ErelGV-AC” isolate was found in infected larvae on cassava plants collected at Cruzeiro do Sul (Acre State, Brazil). The ErelGV-PA isolate was found in infected larvae present on rubber trees collected at Belem (Para State, Brazil) (Table 1). Viral particles were purified according to [20]. In brief, viral-infected dead caterpillars were macerated in homogenization buffer (1% ascorbic acid, 2% SDS, 10mM Tris, pH 7.8, 1mM EDTA, pH 8.0), filtered through cheesecloth layers and centrifuged at 10,000 x *g* for 15 min at 4°C. The pellet was suspended in 10 mL of TE buffer (10mM Tris-HCl, pH 8.0 and 1mM EDTA, pH 8.0) and submitted to another centrifugation step at 12,000 x *g* for 12 min, at 4°C. This new pellet was resuspended in TE buffer and applied onto sucrose gradients with densities varying from 1.17 g/mL to 1.26 g/mL. The gradients were submitted to centrifugation at 100,000 x *g*, for 40 min at 4°C. The granule-containing band was collected, diluted in TE buffer and centrifuged at 12,000 x *g* for 15 min at 4°C. Finally the viral particles (granules) were suspended in water and stored at - 20°C.

**Figure 2.**
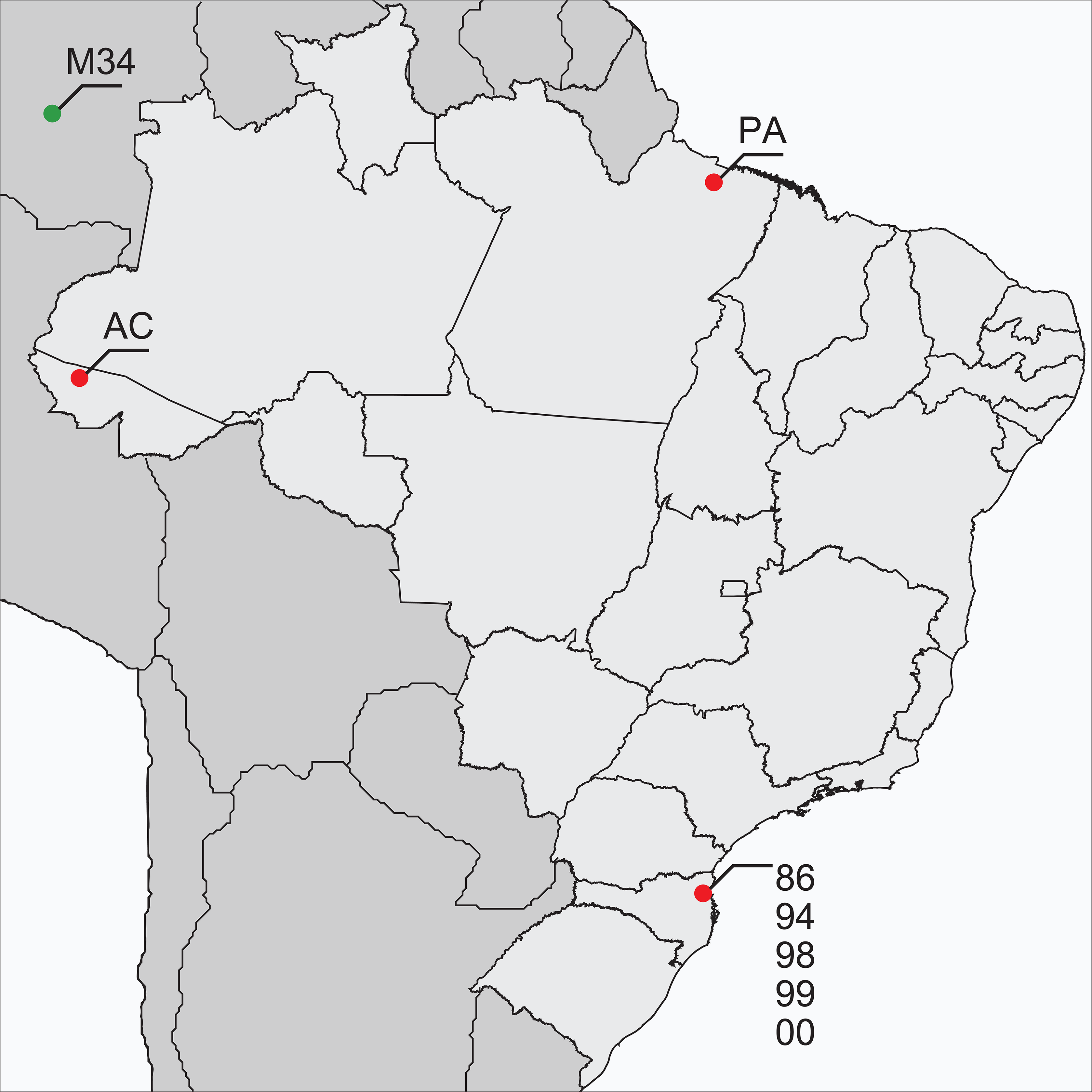
Geographical locations where ErelGV samples were isolated from (approximate coordinates). Red dots depict Brazilian Isolates, *and the green one (*top-left corner) represents a Colombian isolate shown here for illustrative purposes (see phylogenetic section for more details).

### DNA extraction

Granules (1.5 mL) were dissolved by addition of a final concentration of 0.1M sodium carbonate solution and incubation at 37°C, for 30 min. [21]. Viral disruption buffer (10 mM Tris, pH 7.6, 10mM EDTA, pH 8.0, 0.25% SDS) containing 500 μg/mL Proteinase K was added to the sample which was incubated at 37°C (overnight). Viral DNA purification was carried out by extraction cycles of phenol; followed by phenol:chloroform:isoamyl alcohol (25:24:1), and chloroform:isoamyl alcohol (24:1), according to [22]. The DNA was precipitated with absolute ethanol and 3M sodium acetate, pelleted, and washed with 70% ethanol. After air drying, the DNA was suspended in TE buffer and kept at 4°C. DNA quantification was carried out by spectrophotometry using a spectrophotometer Thermo Scientific NanoDrop^TM^ 1000 (220nm – 750nm).

### Genome sequencing, assembly, and sequence analysis

ErelGV isolates were sequenced in a GS FLX Titanium platform (Roche 454 Life Sciences) at the ‘*Centro de Genômica de Alto Desempenho do Distrito Federal*’ (Brasília, Brazil), following the manufacturer’s recommendations (Sequencing Method Manual, GS FLX Titanium Series, Roche 454 Life Sciences). Genome assembly were performed *de novo* using Geneious R7 [23]. A single representative genome was reconstructed from each isolate, considering the following level of stringency: minimum overlap of 150 base pairs among reads, and minimum sequence identity of 97%. Potential out-of-frame sequence errors observed after the assembly were manually inspected and corrected in pairwise comparisons with the genome of ErelGV-86 [11], from which the annotations were transferred into the newly assembled genomes. To analyse the intra-populational diversity, each sequence dataset was mapped against its respective representative genome, and Geneious R7 was used to identify SNPs inside and outside coding sequences of all isolates. After such process, only variants supported by a minimum of five reads and showing at least 1% of frequency were considered, and substitutions within tandem repeats were ignored. In order to assess the diversity of the whole ErelGV metapopulation, a large dataset of over 210,000 reads were created by combining sequences from all viral isolates, and this dataset was mapped against a final ErelGV consensus genome. Prior to SNP detection, errors in homopolymeric regions were identified and corrected using RC454 [24], where low-frequency variants (< 0.01) and those supported for less than 3 reads were removed. To ensure the reliability of our genetic diversity analysis, polymorphic sites identified in coding sequences were manually curated to avoid potential false-positive detections. Gene Ontology information (biological process, molecular function, cellular component) was retrieved from the records of ErelGV-86 available on UniProt [25]. The betabaculovirus maximum likelyhood (ML) tree was inferred in PhyML [26], with 500 replicates under a WAG+I+G model selected by ProtTest [27], using concatenated amino acid sequences of 38 core genes, aligned with MAFFT [28]. ErelGV-specific ML tree was inferred using concatenated partial sequences of *granulin, lef-9* and *lef-8* genes, also aligned with MAFFT, and inferred using the FastTree method implemented in Geneious R7 [23]. The genomic circular map was plotted using Circos [29].

## RESULTS

### General aspects of ErelGV genomes

All isolates of ErelGV included in this study are collinear and no gene gain or loss was observed: all genome encode at least 130 ORFs (minimum size of 150 bp), as observed for ErelGV-86 [11]. When compared to the isolate 86, most genomes showed high nucleotide similarity (from 99.47% to 99.94%) and average G+C content of 39.75% (Table 1). Genome length ranged from 102,616 to 102,764, and insertions/deletions (10 or more base pairs) were found mainly within *direct repeats* (*drs*), *i.e.* tandem repeats located in coding and non-coding regions (Table S1).

By performing whole genome alignments, large indels (12-88 bp) were detected mainly in seven *drs.* Some of them were found within coding regions, leading to size variations in low-complexity regions, as observed in: *erel44* (*dr8*, with three size variants); *p10* (*dr10*, with three size variants); and *erel121* (*dr15/16*, with three size variants). Three other *direct repeats* with large indels were found in non-coding regions located: downstream of *erel11* (*dr2*, showing up to 46-bp indel); downstream of *erel19* (*dr3*, showing up to 26-bp indel); and upstream of *erel23* (*dr5*, showing up to 88-bp indel). No promoter motifs were found in these intergenic *drs*, except for *dr5*, where a putative TATA box motif is disrupted due to deletion in ErelGV-98, -AC, and -PA.

### Intrapopulational genetic diversity

To analyse the intra-isolate diversity, each read dataset from the sequenced isolates was mapped against its respective representative genome, and polymorphisms inside and outside the coding sequences were detected. On average, around 52% of the polymorphisms observed in each isolate corresponded to synonymous substitutions; 38.5% to non-synonymous; and 9.5% to substitutions in non-coding regions. The intra-isolate diversity was slightly similar in most samples, except for ErelGV-AC and ErelGV-PA, which showed extremely high and low levels of diversity, respectively. The total number of polymorphisms ranged from 20 (in ErelGV-PA) to 1,267 (in ErelGV-AC) (Figure 3A), and the number of SNPs observed for each isolate did not correlate to the sequencing coverage (r = 0.015) (Figure 3B).

**Figure 3.**
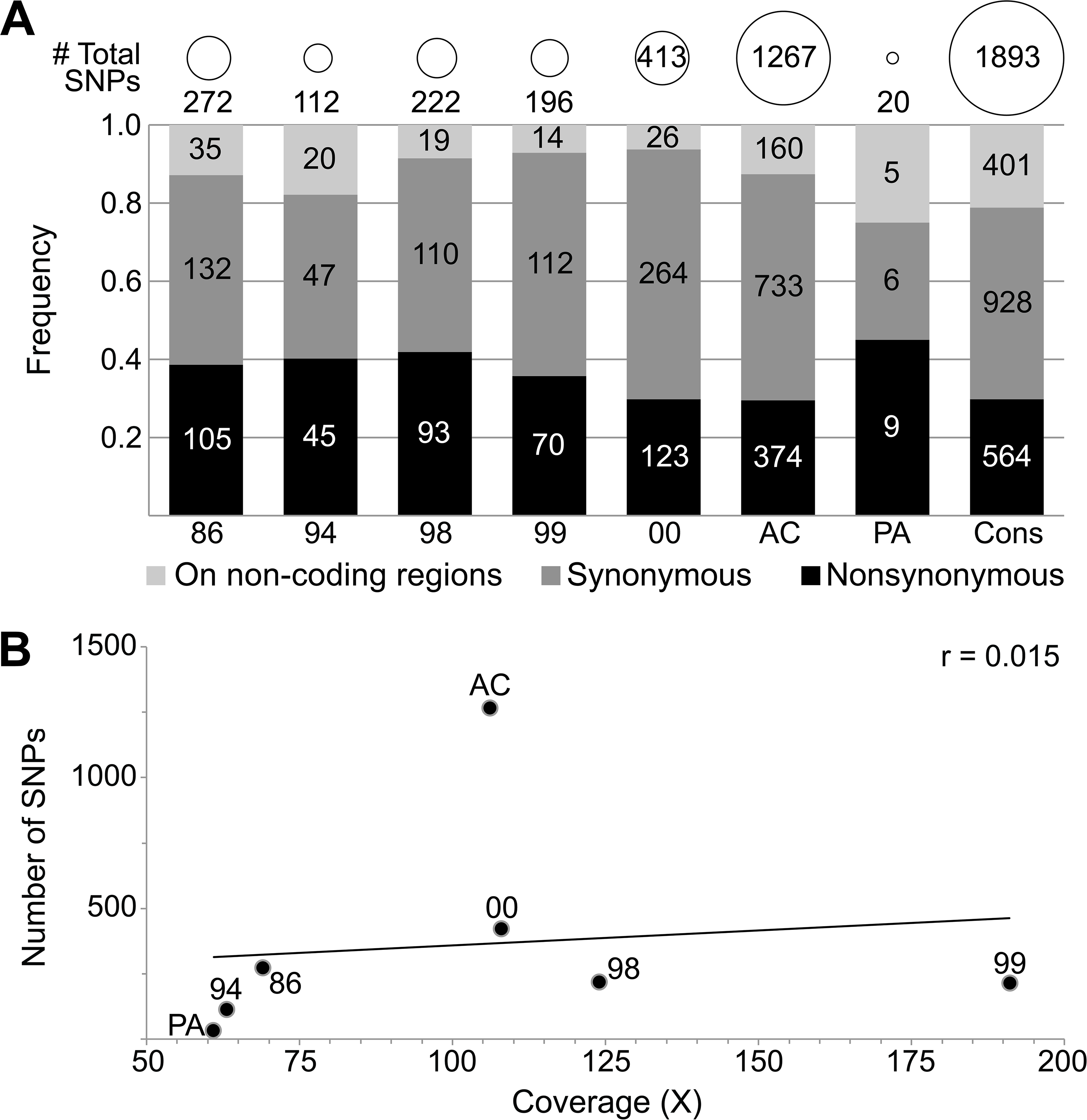
Overview of SNPs on ErelGV genomes. A) Total number of SNPs, and frequency of synonymous, non-synonymous substitutions, and SNPs on non-coding regions of ErelGV isolates. B) Relationship between number of SNPs and sequencing coverage. As observed, these two variables are not correlated (r = 0.015), showing that sequencing depth did not influence the levels of diversity detected in each isolate.

ErelGV ORFs range from 153 to 3,303 bp, and taking into account their differences in size and polymorphisms, genetic diversity was estimated by means of the number of non-synonymous substitutions per base pair (NSS/bp). This approach revealed that most of the genes are either conserved (no NSS) or showed low levels of diversity (from 1 to 3×10^-3^ NSS/bp) (Figure 4 and Figure S1). Moreover, low correlations were observed between diversity and ORF size (-0.01 < *r* < to 0.29). As expected, the exception was ErelGV-AC, which showed moderate correlation for such genetic aspect (r = 0.516), with some genes showing levels of diversity greater than 9×10^-3^ NSS/bp (Figure S1).

**Figure 4.**
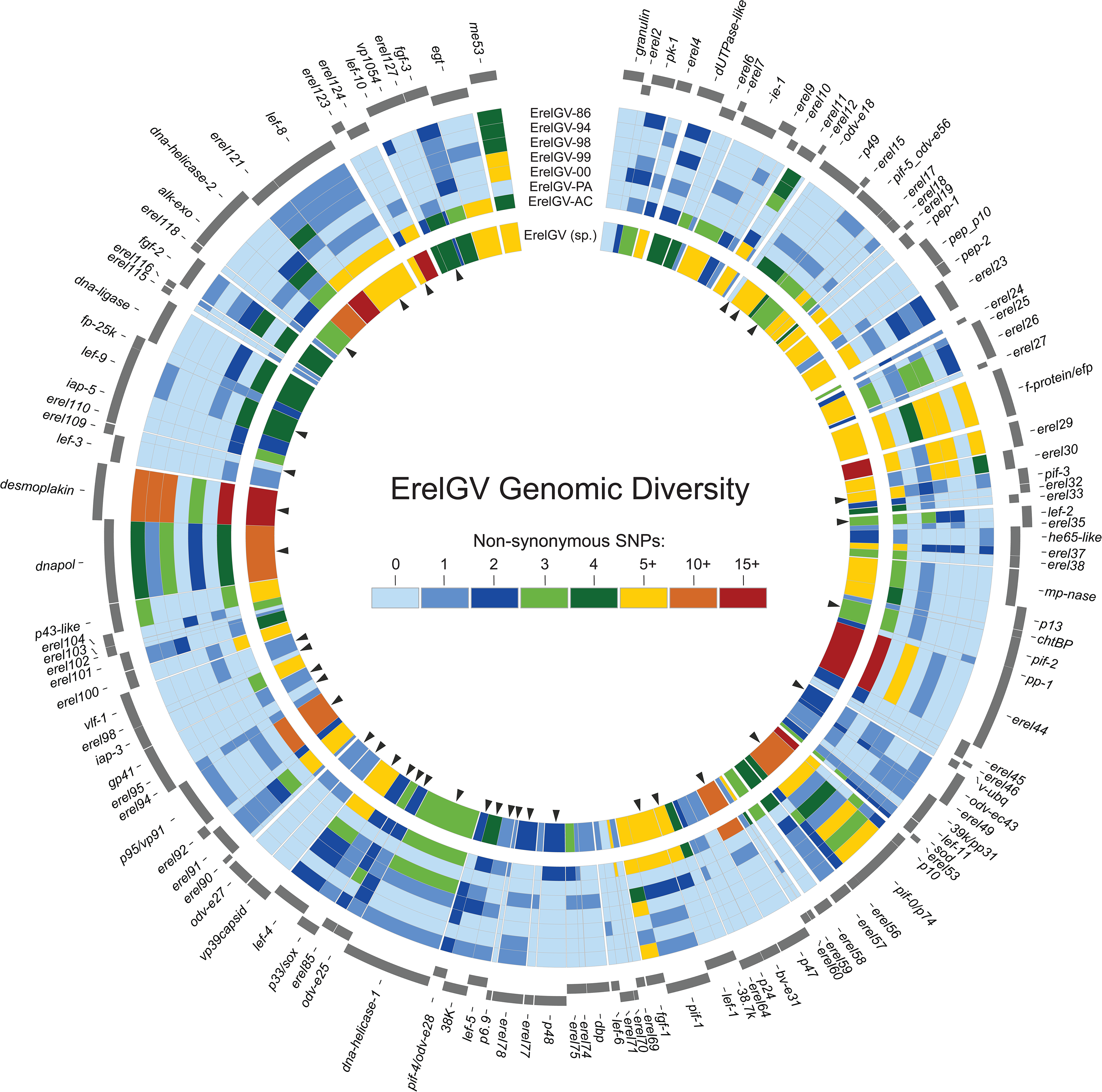
Genetic diversity of the ErelGV. Externally in this circular map ORFs are represented in positive and negative sense. Arrowheads highlight genes shared by all baculoviruses (core genes). From the first to seventh ring the heatmaps depict the number of NSS per gene. Finally, the inner ring summarizes the genetic diversity of ErelGV sp. when all reads are combined and mapped against a consensus genome. See Table S2 for quantitative data.

### Polymorphisms of ErelGV

We have also estimated the diversity in ErelGV species as a species complex. Firstly, we assumed all isolates as members of a hypothetical ErelGV metapopulation, and their reads were combined and mapped against a consensus genome. As SNPs present in frequency and coverage below the minimum thresholds cannot be detected in some isolates, by combining all isolate-specific reads, rare polymorphisms could be identified in the metapopulation due to the cumulative effect provided by all reads in association. Conversely, low frequency isolate-specific SNPs could not be detected when all reads were combined and mapped against the consensus.

The full mapping against the consensus genome revealed at least 1,893 substitutions in coding and non-coding regions. Although around 6% of the ErelGV genome correspond to non-coding regions, more than 21% of the substitutions (401) were located in such regions. Moreover, a total of 564 non-synonymous substitutions were found spread all over the genome, with variable genes interleaved among genes with few polymorphisms (inner ring, Figure 4).

The ErelGV scatterplot of ORF sizes against non-synonymous substitutions has revealed moderate correlation between these two variables (r = 0.514), and the distribution was similar to that observed for ErelGV-AC (Figure 5A and Figure S1). Even after combining all reads, at least 20 invariable genes were identified in the ErelGV species, and seven of them are core genes: *ac78-like* (*erel98*), *vlf-1* (*erel99*), and genes encoding structural proteins like *odv-e18* (*erel13*), *odv-e27* (*erel89*), *p6.9* (*erel79*), *erel95* (*ac81-like*), and *erel109* (*ac68-like*). On the other hand, some genes showed high levels of diversity (more than 15×10^-3^ NSS/bp), as the structural genes *ac53-like* (*erel123*), *ac110-like* (*erel53*), *pif-3, erel17, p10,* and *pif-7*, which is the 38^th^ recently identified core gene [14], and 11 non-core genes with unknown function.

**Figure 5.**
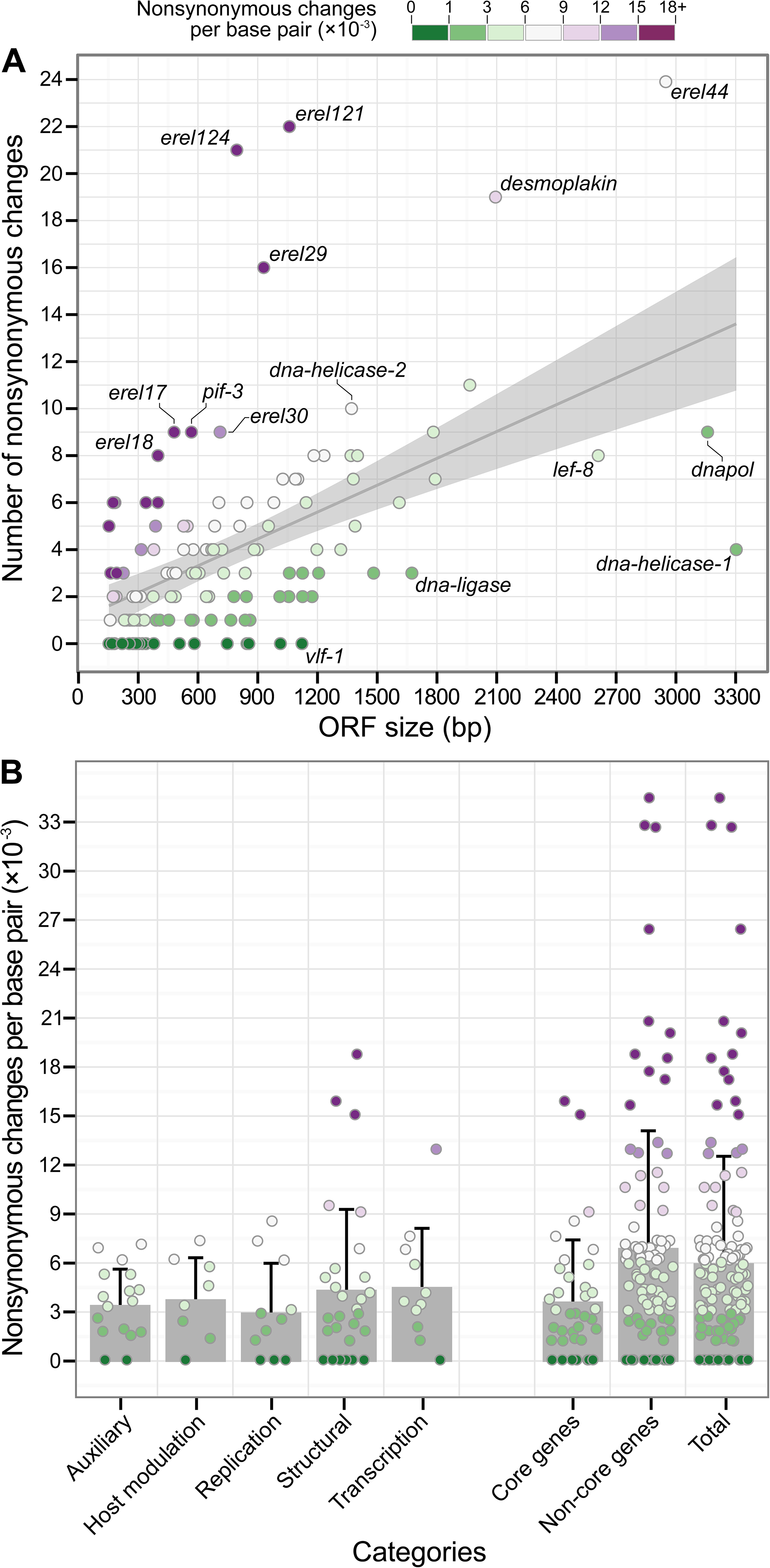
Non-synonymous substitutions, ORF size and functional categories. In these plots, each dot represents a gene depicted in Figure 4. A) Number of NSS per base pair (×10^-3^) and ORF size (bp) have shown moderate correlation (r = 0.51). The grey area corresponds to the 95% confidence interval, and highly conversed or highly diverse genes are shown as labelled outliers. B) Scatter plot showing how genetic diversity (NSS/bp) relates to gene functions. As shown, most highly diverse genes still have unknown functions. For the sake of clarity, the structural gene p10, which shows 53.85 NSS per base pair (×10^-3^), are not included in this plot. See Table S2 for quantitative data.

### Diversity on functional groups

By grouping ErelGV genes based on their main function, we analysed the level of polymorphisms in genes belonging to the following categories: Auxiliary; Host modulation; Replication; Transcription; Structural, as well as core and non-core genes (Figure 5B).

Genes implicated in replication and DNA metabolism, such as *alk-exo, dnapol*, and *dna-helicase-1*, were the most conserved genes in the ErelGV genomes, with an average diversity of 3.03×10^-3^ NSS/bp. Among these genes, *vlf-1, lef-3*, and *dbp* do not show any polymorphisms in all isolates, and, conversely, *lef-1* was the most variable. Genes implicated in host modulation and auxiliary functions have shown similar intermediate levels of polymorphisms, varying on average between 3.49 and 3.84×10^-3^ NSS/bp, respectively (Figure 5B). Genes involved in structural and transcriptional functions were among the most variable ErelGV genes of known function, showing respectively 4.42 and 4.59×10^-3^ NSS/bp of average diversity. Our analyses have revealed that the structural genes *pif-3, desmoplakin, pep-1, erel17*, and *erel123* (*Ac53-like*, a core gene), as well as the transcriptional regulatory gene *lef-10,* are among the most polymorphic genes of ErelGV. Conversely, at least six structural genes (*erel95, erel109, granulin, odv-e18, odv-e27* and *p6.9*), and one gene involved in transcription regulation (*lef-6*) have shown no polymorphisms in all viral isolates.

Interestingly, most genes with high number of polymorphisms (>15×10^-3^ NSS/bp) have unknown functions (Figure 5B). Among them are three genes unique to ErelGV (*erel53*; *erel59*; *erel70*), and other seven only encoded by betabaculoviruses (*erel19*; *erel124*; *erel121*; *erel18*; *erel24*; *erel37* and *erel29*). On the other hand, some genes of unknown function have also shown to be invariable in all isolates, such as: *erel2; erel35*; *erel69*; *erel98* (*Ac78-like,* a core gene); *erel102* (unique to ErelGV); and *erel116*. An important difference in polymorphisms was observed between core and non-core genes. On average, non-core genes have nearly double as many non-synonymous substitutions as core gene (6.05 and 3.70×10^-3^, respectively) (Figure 5B), characteristic already reported in other field-isolated alphabaculoviruses [17].

### Phylogenetic analysis

Our phylogenetic analysis with concatenated protein sequences of core genes revealed the evolutionary relationship of ErelGV isolates and other betabaculoviruses. As expected, all ErelGV isolates clustered together, having ChocGV as their most closely related taxon, as observed by [11]. The analysis also revealed two main clades of betabaculoviruses (Figure 6), which showed a slightly different species composition compared to previous studies [30-32]. Instead of placing PlxyGV and AgseGV as basal taxa in clade A, the topology of the *betabaculovirus* subtree shows them in clade B, which includes CpGV and the ErelGV isolates.

**Figure 6.**
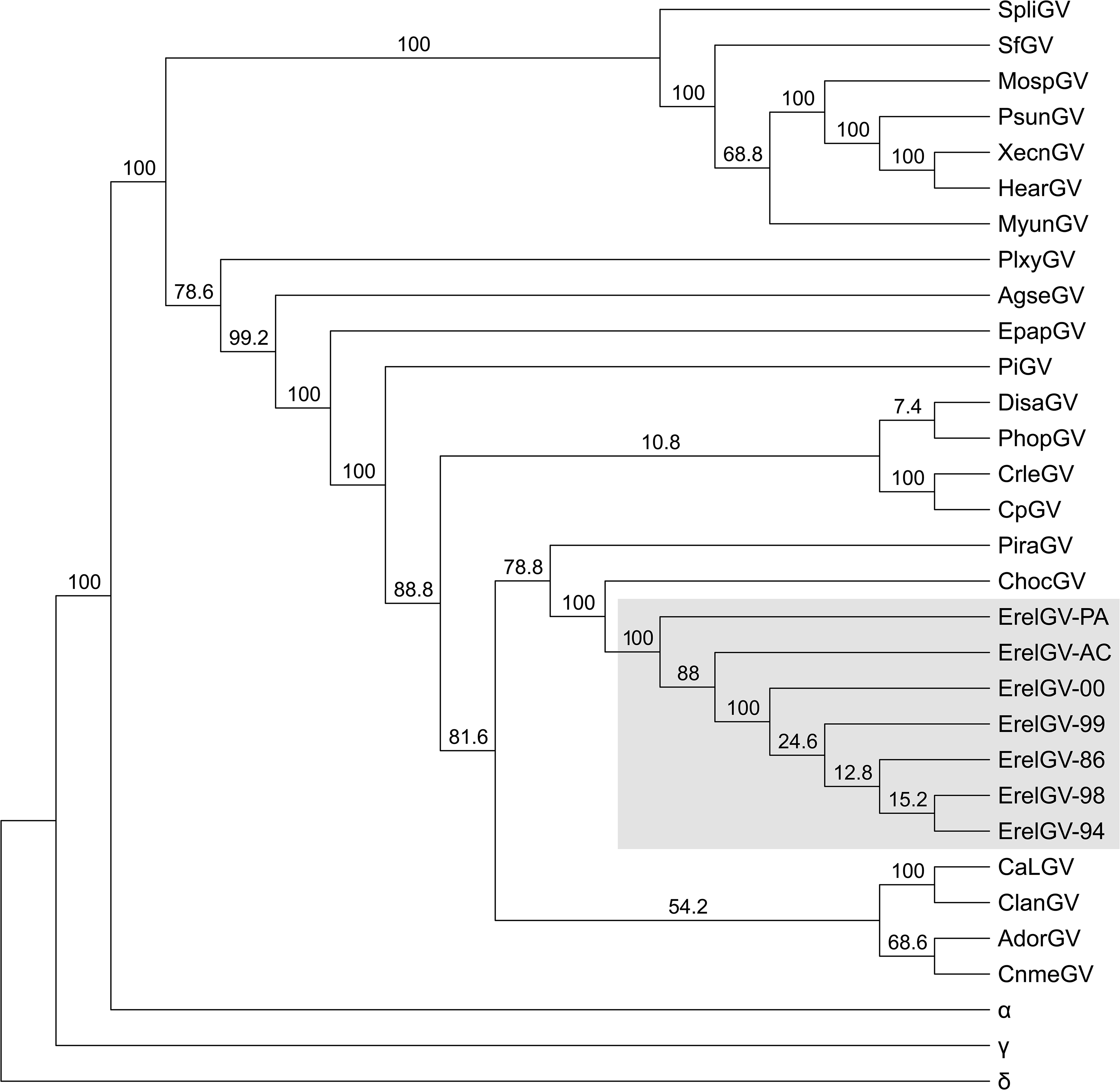
Maximum likelihood tree of *Betabaculovirus* and ErelGV isolates. The phylogeny was inferred using concatenated amino acid sequences of baculovirus core genes. ErelGV isolates are highlighted in grey. The baculoviral genera *Alphabaculovirus* (α = AcMNPV); and *Gammabaculovirus* (γ = NeabNPV); *Deltabaculovirus* (δ = CuniNPV) were included as outgroups, and the latter was used as root. Bootstrap values are indicated for each interior branch. The tree is shown as a cladogram for purposes of clarity only.

In order to better understand the evolutionary history of ErelGV isolates in South America, an additional analysis including the Colombian isolate ErelGV-M34 [33] was performed using concatenated partial sequences of the *granulin, lef-9*, and *lef-8*. ErelGV isolates from the southern Brazilian state of Santa Catarina (SC, see Table 1 and Figure 2) clustered together, having the northern isolates ErelGV-AC and - PA, and the Colombian isolate -M34 as the most distantly related taxa (see Figure S3).

## DISCUSSION

### PIF proteins have distinct evolutionary histories and different sequence diversities

ErelGV encodes at least eight genes known to act as *per os* infection factors (*pif*) (Table 2). A previous study has proposed that proteins P74, PIF1, PIF2 and PIF3 form a conserved complex shared by all baculovirus, which plays essential roles in the baculoviral entry into midgut cells [34, 35]. Our results have revealed that among these genes, the core gene *pif-3* is by far the most variable of them. The protein it encodes shows a conserved hydrophobic (transmembrane) N-terminal sequence [36], and a domain with unknown function (DUF666, e-value: 6.78e-66) at its C-terminal, region where most polymorphisms (6 NSS) were found. Interestingly, this domain is likely to be located externally to the ODV envelope, establishing interactions primarily with other PIFs, but is not directly involved in viral binding or fusion [35]. Furthermore, a recent study identified the core gene p95/vp91 (*pif-8, erel93*) as novel PIF protein required for both ODV and BV nucleocapsid assembly, and the formation of the PIF complex in ODV envelopes [14]. Together with *pif-3, pif-8* and *pif-0* are the three most diverse *pif* genes of ErelGV.

### Paralogous genes evolve under distinct diversifying strategies

Two other putative structural genes were found to be highly variable in this study: *erel17* (18.75×10^-3^ NSS/bp) and *erel123* (15.04×10^-3^ NSS/bp). By homology, the product encoded by *erel17* is suggested to be a viral capsid protein [GO:0019028], and most of the polymorphisms (7 out of 9) were found at its N-terminal region. This region corresponds to a NPV_P10 domain (e-value: 1.40e-07), commonly encoded by the *p10* (*erel54*), and responsible for aggregating P10-like proteins to form filaments and tubular structures [37]. Interestingly, the NPV_P10 domain encoded by *p10* is rather conserved, and its overall gene diversity is much lower than the domain encoded by its potential second copy, *erel17*. Another important difference between the peptides encoded by *p10* and *erel17* lies on their C-terminal regions: while P10 shows a proline-rich region composed by 6-8 copies of a motif “PEPEPESK” (inside *dr10*), EREL17 shows a Serine/Threonine rich C-terminal, apparently not homologous to the one observed in P10. It is unclear what function *erel17* may play, however, since P10-like proteins tend to interact with each other [37], further experimental studies investigating P10-like protein aggregation will be necessary in order to understand their role during host infection.

Another polymorphic structural gene of ErelGV is *pep-1* (9.47×10^-3^ NSS/bp). This gene encodes a protein with a Baculo_PEP_N domain (e-value: 5.1e^-35^), and all *pep-1* NSS were found within this region. Immediately downstream to this gene, two other genes encoding Baculo_PEP_N are found with different levels of diversity, they are: *pep/p10* (6.82×10^-3^ NSS/bp); and *pep-2* (2.21×10^-3^ NSS/bp). The PEP (polyhedron envelope protein) proteins are known to form protective layers that ensure OB integrity in the field [38]. As previously shown [39], alpha-and betabaculoviruses express different numbers of *pep* genes. While alphabaculoviruses encode PEP proteins with Baculo_PEP_N and Baculo_PEP_C domains, the betabaculoviruses show three copies of pep: *pep-1* and *pep-2*, which encode a single Baculo_PEP_N domain; and *pep/p10*, which encodes both the N-and the C-terminal domains. Although PEP protein structures and their binding modes are still unclear [38], the presence of three PEPs with different domain composition and levels of diversity in ErelGV may lead to changes in the OB calyx structure, however, further studies are required to demonstrate it conclusively.

Finally, viral homologs of fibroblast growth factor (*fgf*) gene are shared by both alpha-and betabaculoviruses, and its product (FGF protein) has been suggested to act as a chemoattractant that enhances the migration of uninfected hemocytes towards infected tissues, both promoting viral spread through the larvae circulatory system [40] and accelerating host mortality [41]. Interestingly, all betabaculoviruses encode three *fgf* paralogs (*fgf-1, -2*, and *-3*), which have probably emerged via independent events of duplication or HGT occurring after the betabaculovirus speciation [39]. In ErelGV, *fgf-2* and *-3* have shown similar levels of diversity (3.34 and 4.54×10^-3^ NSS/bp, respectively), while *fgf-1* has shown to be the most diverse copy (7.31×10^-3^ NSS/bp). Interestingly, the ErelGV FGFs have different lengths, and most of their polymorphisms are located within the C-terminal region (Figure S2). FGF proteins are known to contain a signal peptide in its N-terminus, and a C-terminus of variable length [40].

### Some core genes of structural function are highly polymorphic

The *ac53-like* (*erel123*) gene was the second most diverse core gene. This gene is suggested to be involved in both nucleocapsid assembly and transportation [42], and in ErelGV, *erel123* is has a Baculo_RING domain (e-value: 6.04e-44). Interestingly, 5 out 6 polymorphisms found in this gene are located specifically in a Zinc finger motif (IPR013083) of the Baculo_RING domain, however, the impacts of such mutations on gene function are still unclear.

The high level of diversity observed for the structural gene *desmoplakin* (9.07×10^-3^ NSS/bp) coincides with results shown previously, where this gene was pointed as the most variable core gene in baculoviruses [19]. Desmoplakin acts in different viral processes such as: nucleocapsid egression from the nucleus; synthesis of pre-occluded virions; and OB formation [43]. In ErelGV, *desmoplakin* encodes a protein with a conserved Desmo_N domain at its N-terminal, but most NSS (17 out of 18) were found within its C-terminal region, which in turn have no specific domains assigned.

### Genes involved in DNA replication and genome processing: the diversity of LEF proteins

At least three *lef* genes (*lef-1, -2* and *-10*) have shown to be the most diverse genes among those involved in DNA replication and genome processing. LEF-1 and LEF-2 proteins seem to form a complex, where LEF-2 is responsible to bind both the DNA and LEF-1 [44]. All polymorphisms of LEF-1 and LEF-2 were located at their N-terminal regions. While no specific protein domain was found on LEF-1, LEF-2 encodes a single domain (Baculo_LEF-2) that comprises its entire length. The *lef-10* has shown to be the most polymorphic *lef* gene (12.92×10^-3^ NSS/bp). This gene encodes a short protein that is probably a component of a multisubunit RNA polymerase [45]. Although *lef-10* shares 1/3 (134 bp) of its coding region with *vp1054*, only one of its five non-synonymous substitutions is found inside this overlapping region, also causing a synonymous change (CTG→CTA) at the 5’ end of *vp1054*. Since mutations in such regions are likely to impair both genes [46], this pattern evidences an interesting constraint that can limit the adaptive space of some baculovirus overlapping genes.

## CONCLUSION

In this article, we presented a whole genome study on the intra-and inter-isolate diversity of field betabaculovirus populations. ErelGV have been used as an important insect biological control agent on cassava crops, and the present study brings out an extensive analysis on the evolution of this viral species. In terms of phylogenetics, southern Brazil isolates (86, 94, 98, 99 and 00) have shown to be more distantly related to the northern isolates (AC, PA). Our results have also revealed that among SNPs detected by deep sequencing of these isolates, 35-41% correspond to mutations leading to amino acid changes (non-synonymous substitutions). No clear trend was found as to genes gathering into clusters based on diversity, as highly conversed or diverse genes are scattered all over the genome. ORF size and number of NSS were not always proportional, revealing that some genes, especially non-core genes of unknown functions, tend to accumulate more mutations, while others, notably core genes, evolve slowly and gather few or no changes over time. Some genes of ErelGV are present in multiple copies (*p10, fgf, pep*), and each copy shows its particular patterns of evolution. Interestingly, despite theirs names, set of genes known to act in association and/or involved in the same biological processes, as *lefs, pifs* and their respective cognates, are not members of to the same protein family, and also show distinct modes of evolution. More studies are necessary to help us understand the effects of such gene variations on viral infection and fitness.

## DECLARATIONS

### Availability of data and materials

The genomes sequenced in this work are available in GenBank under the accession numbers: KX859079, KX859080, KX859081, KX859082, KX859083, and KX859084.

### Authors’ contributions

Conceived and designed the experiments: MLS, WS; Performed the experiments: MLS, WS; Analyzed the data: AFB, DMPAA, FLM; Contributed reagents/materials/analysis tools: BMR, MLS; Wrote the original manuscript draft: AFB, MLS; Reviewed and edited the manuscript: AFB, BMR, DMPAA, FLM, MLS. All authors read and approved the final manuscript.

## Acknowledgements

We also thank Dr. Renato Arcangelo Pegoraro (EPAGRI-SC), Dr. Murilo Fazolin (Embrapa Acre), and Dr. Orlando Shigueo Ohashi (UFRA) for providing the viral samples. Dr. Murilo Fazolin (Embrapa Acre) for the picture of *E. ello ello* infected by ErelGV. The authors declare that they have no competing interests. The funders had no role in study design, data collection and analysis, decision to publish, or preparation of the manuscript.

